# Drug Target Identification in Triple Negative Breast Cancer Stem Cell Pathways: a computational study of gene regulatory pathways using Boolean networks

**DOI:** 10.1101/2023.05.03.539160

**Authors:** Aditya Lahiri, Haswanth Vundavilli, Madhurima Mondal, Pranabesh Bhattacharjee, Brian Decker, Giuseppe Del Priore, N. Peter Reeves, Aniruddha Datta

## Abstract

Triple-negative breast cancer (TNBC) is an aggressive form of breast cancer associated with an early age of onset, greater propensity towards metastasis, and poorer clinical outcomes. It accounts for 10% to 20% of newly diagnosed breast cancer cases and disproportionately affects individuals from the African American race. While TNBC is sensitive to chemotherapy, it is also prone to relapse. This is because chemotherapy successfully targets the primary TNBC tumor cell but often fails to target the subpopulation of TNBC stem cells. TNBC stem cells display cancerous traits such as cell cycle progression, survival, proliferation, apoptosis inhibition, and epithelial-mesenchymal transition. To study the cancer initiating behavior of the TNBC stem cells, we studied their underlying signaling pathways using Boolean networks(BN). BNs are effective in capturing the causal interactions taking place in signaling pathways. We built the BN from the pathway literature and used it to evaluate the efficacies of eleven targeted inhibitory drugs in suppressing cancer-promoting genes. We simulated the BN when the pathways had single or multiple mutations, with a maximum of three mutations at a time. Our findings indicated that *STAT3, GLI*, and *NF-κB* are the most optimal targets for inhibition. These genes are known regulators of the cancer-promoting genes in the pathway,hence our model agrees with the existing biological literature. Therefore inhibiting these three genes has the potential to prevent TNBC relapse. Additionally, our studies found that drug efficacies decreased as mutations increased in the pathway. Furthermore, we noticed that combinations of drugs performed better than single drugs.

## I. INTRODUCTION

Cancer is the second leading cause of death in the United States and is a major barrier to increasing life expectancy around the world [1]–[3]. Breast cancer is now the most frequently diagnosed cancer and is the leading cause of cancer deaths among women around the world [2], [4]–[7].

It is estimated that there will be 43,780 deaths due to breast cancer in the United States in 2022 [1]. Breast cancer is not a single disease; instead it is an umbrella term for diseases characterized by heterogeneous groups of neoplasms originating from the epithelial cells lining the mammary ducts of the breast [8]–[10]. Historically breast cancer heterogeneity has been studied through clinicopathological features such as tumor stage, grade, histology type, and proliferation status [11]–[13]. With the accumulation of knowledge in breast cancer and the advancement in sequencing technology over the years, it has become evident that tumor characteristics and their response to drugs is governed by their underlying biology [12], [13]. Consequently, breast cancer is now classified based on tumor molecular biology, which provides insights into both the disease mechanism and clinical outcomes [9]. Typically molecular classification of breast cancer is based on the expression levels of well established tumor markers such as estrogen receptor (ER), progesterone receptor (PR), and over-expression and/or amplification of the human epidermal growth factor receptor 2 (*HER2*) [11], [14]. Molecular classification has been used to determine the prognosis and response to endocrine treatment and anti-*HER2* agents.

In this article, we focus on triple negative breast cancer (TNBC), which is characterized by a lack of or low expression levels of ERs, PRs, and *HER2* [11], [14], [15]. TNBC accounts for 10%-20% of newly diagnosed breast cancer cases and compared to other breast cancer subtypes, it is associated with highly aggressive clinical behavior, early age of onset, African American race, higher grade and mitotic index, greater propensity towards metastasis and poorer clinical outcomes due to higher relapse and survival rates [14]–[18]. Since TNBC lacks ER, PR and *HER2*, it does not respond to endocrine treatment or anti-*HER2* agents [19]. TNBC is sensitive to chemotherapeutic drugs such as anthracycline, alkylating agents, taxanes, and antimetabolite fluorouracil, due to which chemotherapy is the standard of care for this phenotype of breast cancer [15], [19]–[21]. However, for patients with relapsed TNBC, there is no standard chemotherapeutic regimen [20].The molecular mechanism underlying the relapse of TNBC is yet not well understood, and no targeted therapeutic agents have demonstrated significant improvement in patient survival, due to this reason, chemotherapy remains the standard of care for TNBC [15]. With chemotherapy, the median progression-free survival ranges from 1.7 to 3.7 months [20], whereas the median overall survival with the onset of metastasis is 23 months [22]. Therefore in this article, we study the underlying biological mechanisms responsible for the relapse of TNBC, specifically, we focus our attention on understanding the biological pathways in TNBC stem cells (TNBCSC). In malignant tumors, cancer stem cells (CSC) such as TNBCSC are a small population of cancer cells alongside the primary tumor cells. Human breast cancer stem cells are associated with tumor initiation, proliferation, metastasis, and resistance to chemotherapy and radiation [23]–[25]. While chemotherapy successfully targets the primary tumor cells, it often fails to target and eliminate the CSCs, leading to relapse [26]. Thus understanding the biological pathways in TNBCSC and identifying druggable targets in those pathways can supplement chemotherapy in the treatment of TNBC and prevent relapse. In the following sections, we will discuss the pathways associated with TNBCSC and then model these pathways using Boolean networks to identify potential drug targets.

## II. SIGNALING PATHWAYS IN TNBCSC

Cells perform critical functions such as metabolism and differentiation through signaling pathways. Signaling pathways consist of molecules such as proteins, genes, and transcription factors which work together in a well-orchestrated manner to perform specific cellular functions. These pathways are activated in the presence of a stimuli or signaling molecule. Signaling molecules such as hormones or growth factors bind to specialized proteins called receptors on the cell surface. The receptors in turn activate their downstream molecules, which initiate a cascade of intercellular signaling activities [27]. The signaling continues till the last molecule in the pathway has been activated. Breakdown in the signaling pathway or abnormal activation of signaling pathways due to genetic mutations can cause aberrant signaling behavior and ultimately lead to the loss of cell cycle control which can cause cancers. Genetic mutations in pathways have been extensively studied and shown to cause cancer [28]–[34]. Genetic mutations are either acquired over the lifetime of the patient or inherited through mutations in the mother’s egg cell or the father’s sperm cells. There are many different types of genetic mutations, and they can be classified broadly as point mutation, chromosomal mutation, and copy number variation [35]. Point mutation refers to changes in a single or a few base-pairs of the genome, these include substitutions, point mutations are also known as single nucleotide variant (SNV) [36]. Chromosomal mutations on the other hand involve inversion, deletion, translocation or duplication of regions of the chromosome (typically spanning longer than one gene) [35], [37]. Copy number variation (CNV) refers to the phenomena where the number of copies of a specific segment of the DNA varies among the genomes of different individuals [35]. In addition to mutations affecting genes, epigenetic factors such as methylation can also affect gene expression in cancer [38]. While there are several types of mutations that affect gene expression, in this study, we focus on the effect of mutations on cancer-promoting activities of the TNBCSC pathways. Our objective is to take a systems-level approach to analyze the effect of mutations on the entire signaling pathways instead of characterizing the nature of genetic mutations. A composite image of the signaling pathways relevant to our study is displayed in Fig.1, and a brief description of each follows.

**FIGURE 1:**
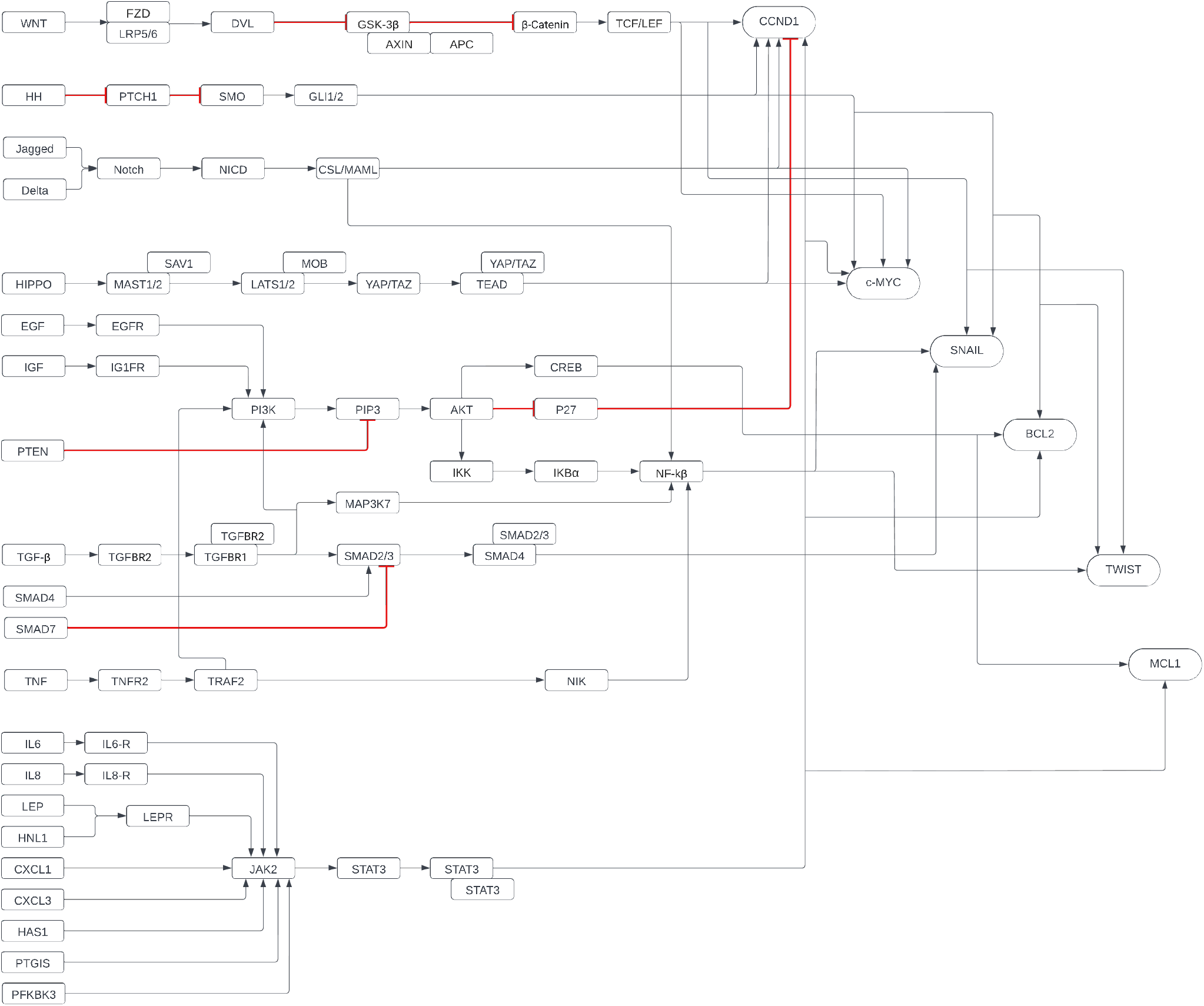
Signaling pathways in TNBCSC.

### A. WNT/β-CATENIN SIGNALING PATHWAY

The *WNT*/*β*-Catenin signaling pathway regulates cell proliferation, migration, and differentiation and has been reported to be highly activated in TNBC [39]–[41]. Furthermore, the *WNT*/*β*-Catenin signaling pathway promotes self renewal and differentiation in breast cancer stem cells [42], [43]. This pathway is activated when the *WNT* ligand binds to the Frizzled (*FZD*) receptor protein and the low-density lipoprotein receptor-related protein5/6 (*LRP*5/6) to form a ternary complex [39], [44]. The activation of the pathway leads to the recruitment of cytosolic (Dvl) protein which inhibits glycogen synthase kinase-3*β* (*GSK*3*β*)/APC/AXIN complex which in turn stops the degradation of*β*-catenin and leads to the accumulation of cytosolic *β*-Catenin. [45]. *β*-catenin further translocates into the nucleus activating T-cell factor/lymphoid enhancing factor (*TCF*/*LEF*) which can regulate genes such as *CCND1, c-MYC* which lead to cell cycle progression, proliferation and the genes *SNAIL* and *TWIST* which code for epithelial-mesenchymal transition (EMT) that can lead to metastases [39], [46], [47].

### B. HEDGEHOG SIGNALING PATHWAY

Several studies have shown the role of Hedgehog (*HH*) signaling pathway in promoting TNBC and breast cancer stem cell renewal, reprogramming, stemness, proliferation, and EMT [48]–[53].The canonical Hedghehog pathway is initiated in the presence of Hedgehog ligands such as sonic Hedgehog, desert Hedgehog or Indian Hedgehog [51]. These ligands bind and inhibit the transmembrane receptor *PTCH*1, which releases transmembrane transducer *SMO* and allows its translocation to the primary cilium. The translocated *SMO* activates the protein *GLI*, which regulates the expression of genes coding for cell cycle progression (*CCND1*), proliferation (*c-MYC*), apoptosis inhibition (*BCL2*) and EMT (*SNAIL*, and *TWIST*) [53]–[58].

### C. NOTCH SIGNALING PATHWAY

The aberrant activation of the *Notch* signaling pathway has been implicated in TNBC progression and breast cancer stem cell renewal, chemotherapy, and radiation therapy resistance [59]–[64]. The *Notch* signaling is activated when transmember ligands *Delta* and *Jagged* (*JAG*) bind with the *Notch* receptors on the cell surface [64], [65]. Upon binding, prote-olytic cleavages in the *Notch* receptors cause the *γ*-secretase-dependent release of the *Notch* intracellular domain (*NICD*) into the cytoplasm [64]. *NICD* translocates into the nucleus and forms a complex containing BF1/Suppressor of Hairless, and Longevity-Assurance Gene-1 (*CSL*), mastermind-like protein (*MAML*), Ski-interacting protein as a CBF1 binding protein (SKIP) and p300 [66]. This complex activates target genes of *Notch* pathway such as *CCND1, c-MYC*, and *NF-κB* [67]–[69].

### D. HIPPO SIGNALING PATHWAY

The *Hippo* signaling pathway plays a critical role in regulating cell proliferation and cell death, which is why aberrant signaling of this pathway has been associated with carcinogenesis and TNBC progression [70]–[73]. Furthermore, the *Hippo* signaling pathway is responsible for self renewal, maintenance, and tumor initiation of breast cancer stem cells [64], [74], [75]. This pathway comprises of mammalian STE20-like kinase 1/2 (*MST*1/2), large tumor suppressor 1/2 (*LATS*1/2), WW domain of the *SAV* family containing protein 1 (*SAV*1), *MOB* kinase activator 1 (*MOB*1), Yes-associated protein (*YAP*) or transcriptional coactivator with PDZ-binding motif (*TAZ*), and members of the TEA domain (*TEAD*) family [64], [70], [76]–[78]. When this pathway is initiated, it phosphorylates *MST*1/2 which activates *LATS*1/2 and leads to the inactivation *YAP*/*TAZ* [75], [76]. However, when this pathway is not turned on, the transcriptional coactivator *YAP*/*TAZ* translocates to the nucleus and induces transcription factor *TEAD* and its target genes, such as *CCND1* and *c-MYC*, which regulate cell cycle and proliferation [79]– [81]. Thus in its deactivated state, the *Hippo* pathway promotes tumor progression.

### E. PI3K-AKT SIGNALING PATHWAY

The *PI3K*(phosphoinositide 3-kinase)-*AKT*(also known as protein kinase B) pathway controls cell proliferation, survival, motility, and differentiation [82]. This pathway is often mutated in mesenchymal TNBC, which overexpresses EMT and breast cancer stem cell features [43], [83]–[86]. This pathway is initiated by receptor tyrosine kinase (RTK) such as epidermal growth factor receptor (*EGFR*), insulin-like growth factor receptor (*IGFR*), and transforming growth factor receptor-*β* (*TGFR*-*β*) etc [87]–[89]. RTKs activate *PI3K*, which leads to the downstream phosphorylation of AKT through phosphatidylinositol-3,4,5-triphosphate (*PIP3*) [90], [91]. The pathway is also regulated by tensin homolog (*PTEN*), which inhibits this pathway and functions as tumor suppressor [91], [92]. About 25%-30% of TNBC cases are characterized by *PTEN* loss mutation, which leads to the aberrant signaling of the *PI3K*-*AKT* pathway [91]– [94]. When this pathway is activated, phosphorylated *AKT* regulates downstream molecules such as cAMP response element-binding protein (*CREB*), *p27*, and *NF-κB* (via the kinase subunit *IKKα*) [75], [95].This pathway regulates the apoptosis inhibitor and survival genes *BCL2, MCL1* (respectively) through *CREB*, inhibits the cell cycle progression gene *CCND1* though *p27*, and the EMT genes *SNAIL* and *TWIST* via *NF-κB* [96]–[100].

### F. TNF SIGNALING PATHWAY

The Tumor Necrosis Factor (*TNF*) signaling pathway is initiated by the pro-inflamtory cytokine *TNF*-*α* which is involved in cancer progression, including that of TNBC [101], [102]. High *TNF*-*α* expression has been linked with increased TNBCSCs via upregulation of *TAZ* and *NF-κB* [102]. Additionally, two studies indicated that the EMT genes *TWIST* and *SNAIL* are upregulated with increased *TNF*-*α* levels [103], [104]. A study by Blakwill showed that increased *TNF*-*α* induces EMT and stemness via the *NF-κB* [105]. *TNF*-*α* activates this pathway by binding to the receptor *TNFR2*; *TRAF2* weakly binds to *TNFR2*, which activates *NF-κB* through *NF-κB*-inducing kinase (*NIK*) [106]. This leads to the activation of EMT genes *SNAIL* and *TWIST*.

### G. TGFβ SIGNALING PATHWAY

The Transforming Growth Factor(TGF)*β* signaling pathway is complex and works as a double edged sword in cancer progression. This pathway promotes apoptosis and cell cycle arrest in early stage cancer cells, however, in later stages, it enhances metastases, invasion, EMT, and drug resistance [107]–[109]. In TNBC, activation of the *TGF-β* pathway in cancer stem cells has been implicated in advancing drug resistance [75], [108], [110]. The canonical *TGFβ* pathway is activated by the binding of the *TGFβ* ligand to the receptor *TGFBR2*, which then further complexes with *TGFBR1. TGFBR1* phosphorylates *SMAD2/3* which leads to a complex formation with *SMAD4* [111]. This complex translocates to the nucleus, where it activates the EMT gene *SNAIL* [111]. *SMAD7* inhibits the *TGFβ* canonical signaling by binding to *SMAD2/3* [111], [112]. The other EMT gene *TWIST*, is activated through non canonical *TGFβ* pathway where *TGFBR1* activates NF-*κ*B signaling via *MAP3K7* [111]. As mentioned earlier, this pathway also activates *PI3K* signaling, and this is mediated through *TGFBR1*.

### H. JAK-STAT SIGNALING PATHWAY

The *JAK*-*STAT* pathway regulates cell proliferation, motility, and stemness [113]. The *JAK*-*STAT* pathway can be activated by various growth factors, cytokines, ligands, and genes that bind to a *JAK* receptor. In TNBC, this pathway can be activated through interleukin(*IL6*), *IL8*, (*LEP*) Prostaglandin-I synthase (*PTGIS*), Hyaluronan synthase 1 (HAS1), C-X-C Motif Chemokine Ligand 3 (*CXCL*3), and 6-phosphofructo-2-kinase/fructose-2, 6-biphosphatase 3 (*PFKFB3*) [75], [114]–[117]. Furthermore, Marotta et al. showed that the *IL6*/*JAK2*/*STAT3* pathway is highly activated in TNBCSC with an increased risk of metastasis compared to other breast cancer stem cells [114]. Activation of a *JAK* receptor by one of the above mentioned growth factors, cytokines, ligands, or genes can lead to the phosphorylation and dimerization of *STAT3* [118]. *STAT3* translocates to the nucleus and activates genes coding for proliferation, survival such as *c-MYC, CCND1, MCL1*, and EMT such as *TWIST* and *SNAIL* via cross talk with NK-*κ*B [119]–[124].

## III. METHODS

The signaling pathways we studied above tightly regulate various aspects of the cell, such as proliferation, survival, apoptosis, and EMT. Under normal conditions, these pathways are tightly regulated to maintain the natural homeostasis of the cell. However, genetic mutations in these pathways lead to loss of cell cycle control and cause diseases such as cancer. Therefore, studying these pathways can reveal further insights into the dynamics of these pathways and help discover the potential drug targets. Computational models have been extensively used in the past to study biological signaling pathways [125]–[130]. Signaling pathways have been successfully studied using various computational methods such as linear models, differential equations, Boolean networks, and Bayesian networks [131]–[133], [133]–[136].The complex interactions taking place in signaling pathways describe a cause-effect relationship between an upstream and downstream molecule. In order to model such interactions, we use Boolean Networks (BN). BNs integrate pathway information from biological literature, and also allows us to study the effects of mutations and drug intervention in the pathways, which is key to identifying the best targets in the pathway for drug intervention. Furthermore, due to the lack of publicly available large scale gene expression data pertaining to TNBCSC, we cannot reliably employ data-driven models such to analyze these pathways. BNs are deterministic in nature and do not require data for modeling, thus BNs helps overcome the hurdle of lack of data availability. Thus BNs serve as an appropriate choice of modeling technique for studying TNBCSC pathway. We use the BN model to study the pathway functioning under normal (healthy) conditions and in the presence of genetic mutations. Furthermore, we augment the model to evaluate the effect of small molecule inhibitor drugs on genetic mutations and rank the drugs according to their efficacy. In the following section, we will describe the basic concept involved in creating a BN.

### A. MODELING SIGNALING PATHWAYS WITH BOOLEAN NETWORKS

Modeling gene regulatory networks using BN was first described by Stuart A. Kauffman in 1969 [137]–[139]. A BN consist of a set of binary valued variables X ={x_1_, x_2_, x_3_,..,x_*n*_} and a vector of Boolean functions F ={f_1_, f_2_, f_3_,..,f_*n*_} [140]. BN can also be viewed as a directed graph G, where the elements of the set X represent the nodes or system variables, and the nodes are connected to each other through directed edges. The nodes are binary valued so they can be active (1) or inactive (0). The causal interactions occurring among the nodes is described by the vector of Boolean functions F [141]. In the context of signaling pathways, the elements of the pathways such as gene and transcription factors can either be upregulated (activated) or downregulated (inhibited); this can be modeled using the binary framework of Boolean Networks. We illustrate BN modeling through a toy signaling pathway in Fig.2.

**FIGURE 2:**
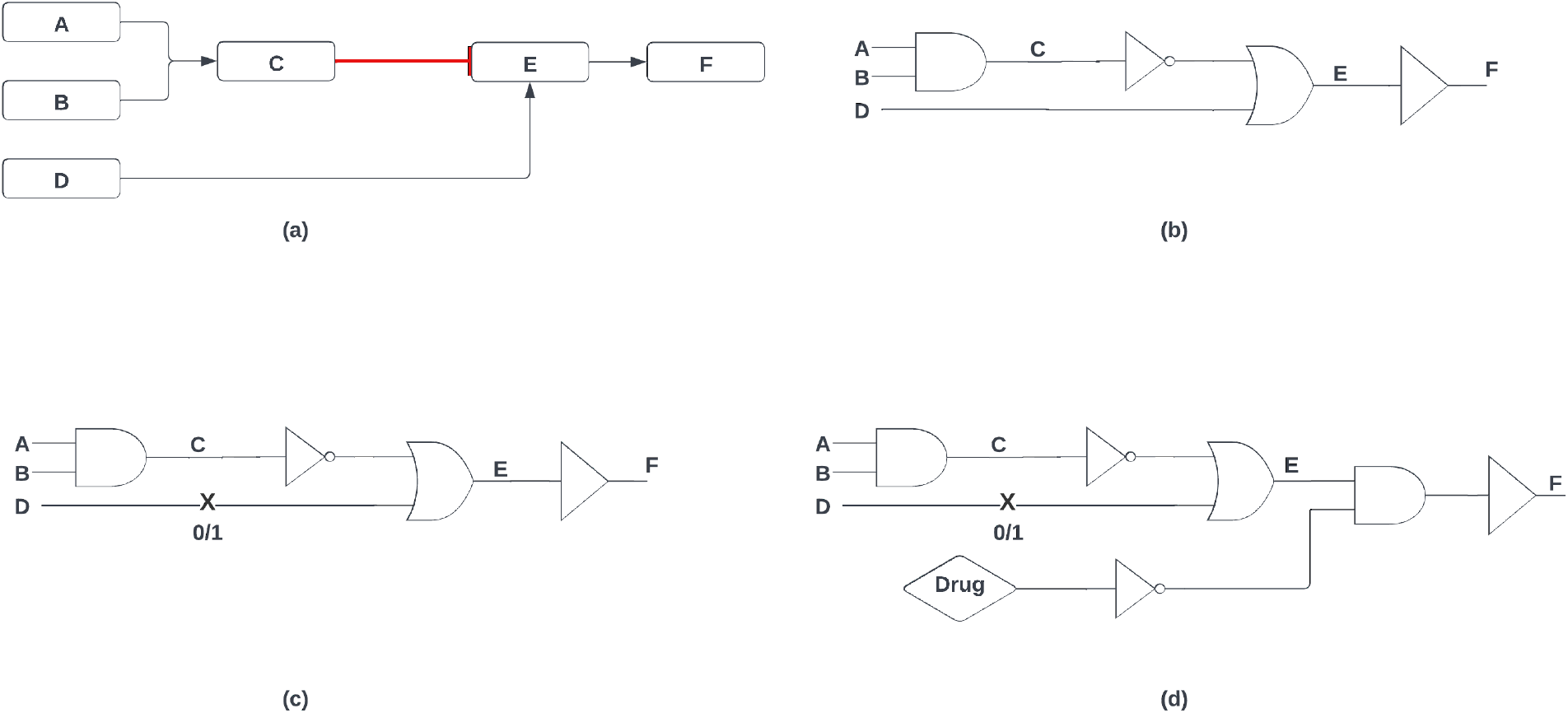
(a) Example signaling pathways. (b)Boolean network model of example signaling pathway. (c) Boolean network with fault at gene D. (d) Boolean network with drug intervention at F

In Fig.2(a), we have a signaling pathway consisting of 6 genes (A-F). When genes A and B are upregulated, they can simultaneously activate gene C. Activated gene C inhibits gene E, whereas an upregulated gene D activates gene E. Activated gene E can upregulate gene F. In this example, we consider that these genes can be only activated (upregulated) or inhibited (downregulated) to model them using BN. Fig.2(b) represents the Boolean network equivalent of the pathway depicted in Fig.2(a). Since genes A and B both need to be activated to turn on gene C, this relationship can be modeled using the logical AND gate, the equivalent Boolean function representing this relation is given by C= A ∧ B. The gene C suppresses gene E, this is why the signal from gene C to gene E first passes through a NOT gate. This signal then feeds into an OR gate along with the input signal from gene D to regulate gene E. Thus, the Boolean function for gene E is given by: E= (¬C) ∨. Gene E activates gene F, and this interaction is represented using a follower or buffer, and the Boolean function for gene F is given by: F=E.

### B. MODELING GENETIC MUTATIONS WITH BOOLEAN NETWORKS

In the earlier sections, we discussed that genetic mutations in the pathways can lead to aberrant signaling which can ultimately lead to cancer. Thus we need a method to model genetic mutations in BNs to model TNBCSC signaling pathways. In the presence of a genetic mutation in a signaling pathway, the mutated element is either silenced or hyper activated. This behavior due to mutations can be modeled as a “stuck-at fault” in BNs. When an element of a BN is stuck-at fault, its state is fixed at either 0 (inhibited) or 1 (active), and can no longer be influenced by inputs from other nodes in the network. Fig.2(c) represents the BN equivalent of the pathway in Fig.2(a) with a stuck-at-fault at gene D. If gene D is stuck-at the state of 0 (deactivated), then its input to gene E will always be 0, thus for gene E to be active, it will require genes A and B should not be simultaneously active(1). Similarly, if gene D is stuck-at the state of 1 (active), Gene E will always be active (1), regardless of the states of gene A or B, as gene E is related to gene D through an OR gate, which requires only one active (1) input to produce an active (1) output.

### C. MODELING DRUG INTERVENTION WITH BOOLEAN NETWORKS

Drugs enhance or suppress the activity of their target molecule by binding to them. This interaction can be modeled in BN by forcibly suppressing or enhancing the target molecule of the selected drug. Fig.2(D) demonstrates the application of a drug that inhibits its target molecule gene F. This interaction between the drug and gene F is mapped in through a NOT gate followed by an AND gate in the BN. In the presence of the drug (state 1), it sends an inhibitory signal (0) to gene F. Gene F also receives a signal from gene E, thus both gene E and drug influence gene F, and this interaction is represented using the AND gate. Since the AND gate receives an inhibitory signal (0) from the drug, it will successfully inhibit gene F. It should be noted that the mutation in gene D which is represented by a stuck-at fault is no longer able to influence the state of gene F in presence of the drug.

In this study, we evaluate the effect of eleven different small molecule inhibitory drug classes on a TNBCSC signaling pathway with faults (mutations). A total of 42 unique faults locations were considered. The objective was to determine how effective these drugs were in mitigating the aberrant behavior caused by faults in the TNBCSC network. Table 1 outlines the eleven drug classes and their molecular targets; we only consider drug classes and not specific drugs as this study is computational in nature, and the choice of drugs is left to clinicians. Our study focuses on identifying the best drug targets; therefore we are only concerned with the drug classes. Furthermore, for some drug targets, there might not be currently approved drugs in the market, and the approach used in the current study can reveal the potential value of designing drugs for these targets.

**TABLE 1:**
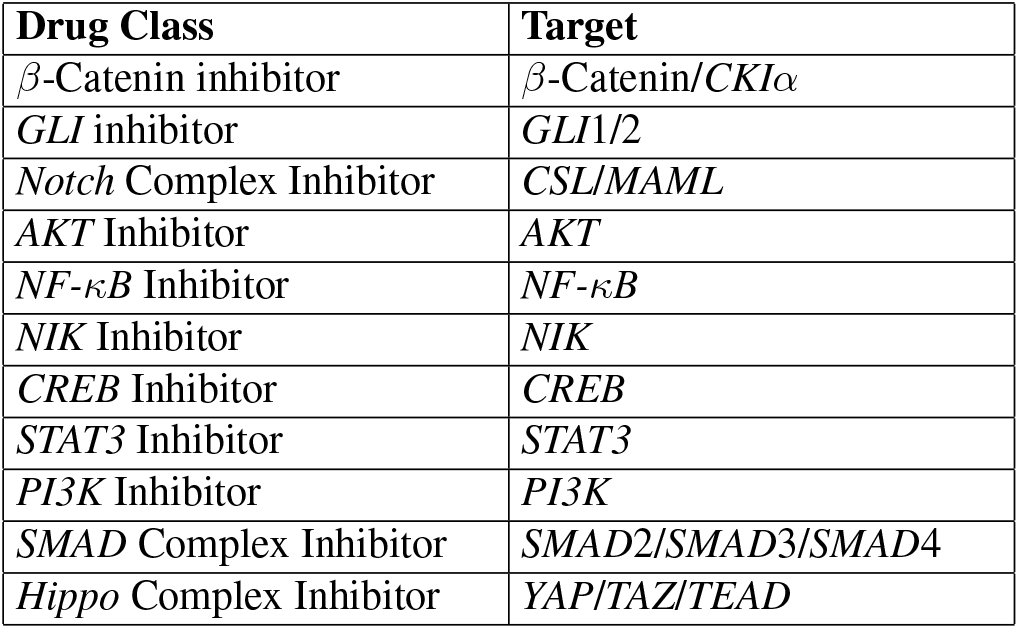
Drug Classes and Targets

We restrict our drug choices to mostly downstream targets as they will be effective in countering aberrant signals originating from upstream faults in the network. The Boolean network model of the TNBCSC pathways (Fig.1) is displayed in Fig.3. Each of the 42 fault locations are numbered beneath each network component. Faults within parenthesis are stuck-at-1, while stuck-at-0 faults are denoted within square brackets. It should be noted that in Fig.3 only the fault sites are shown, but none of the faults are activated.

**FIGURE 3:**
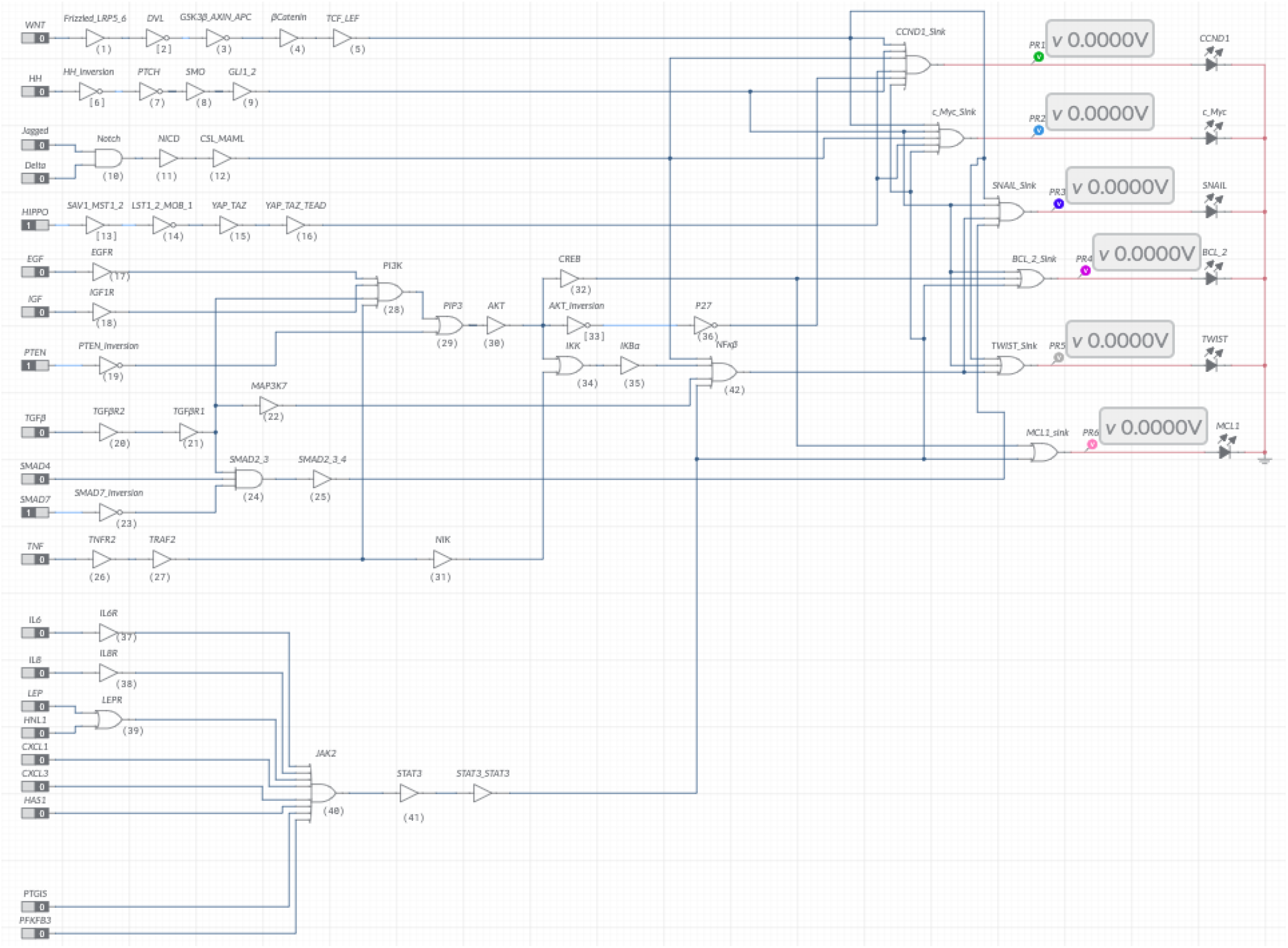
Boolean Network model of signaling pathways in THBCSC.

### D. SIMULATION OF TNBCSC PATHWAYS

From a system level view, the BN representing the TNBCSC can be viewed as a multi-input multi-output (MIMO) digital circuit. In the absence of faults, the system output is governed solely by its inputs. Thus a fault-free network is representative of the pathways of a healthy cell. On the other hand, for a given input, a BN with faults will have a different output than a fault-free BN, describing the aberrant signaling in cancerous cells. Therefore, a cancerous cell will not have the same input-output mapping as that of a healthy cell. For the BN network considered in this study, there are 21 inputs consisting of growth factors, pathway receptors, interleukins, and tumor suppressors, whereas the five outputs consist of cancer promoting genes. Thus for a healthy cell, a healthy input state means that the tumor suppressors *PTEN,SMAD*7, and *Hippo* are turned on (state 1), the rest of the inputs are turned off (state 0). Furthermore, a healthy input state also implies that all the cancer promoting genes at the output of the BN are turned off. This is indeed the case, and is reflected in the fault-free network in Fig.3, where the inputs are in a healthy state and the outputs genes are all turned off, as indicated by the zero reading at each of the cancer promoting genes at the output of the BN. The healthy input and its corresponding output for a fault free BN depicted in Fig.3 is defined as follows in Table 2 and Table 3 respectively.

**TABLE 2:**
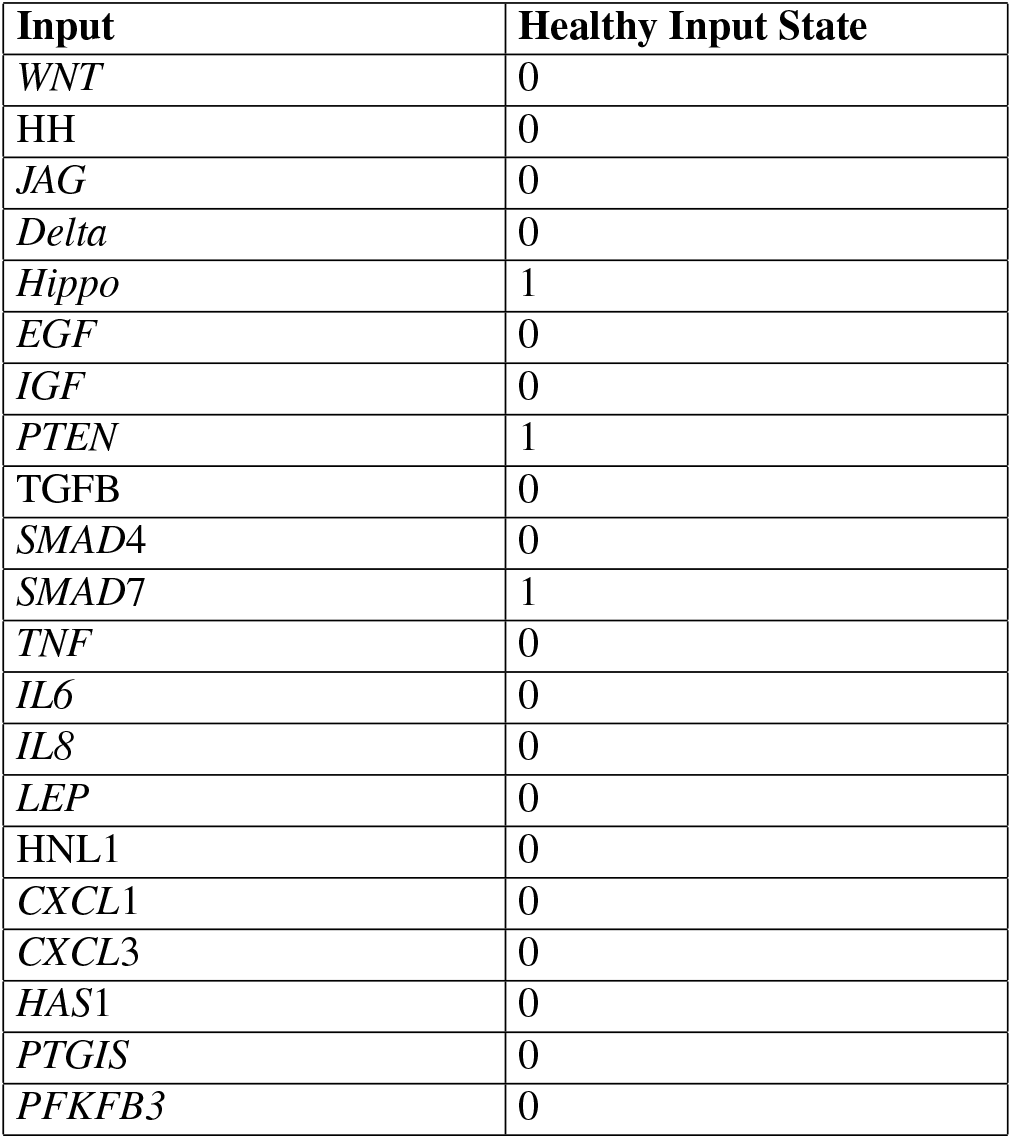
Boolean network (fault free) inputs and their corresponding healthy state

**TABLE 3:**
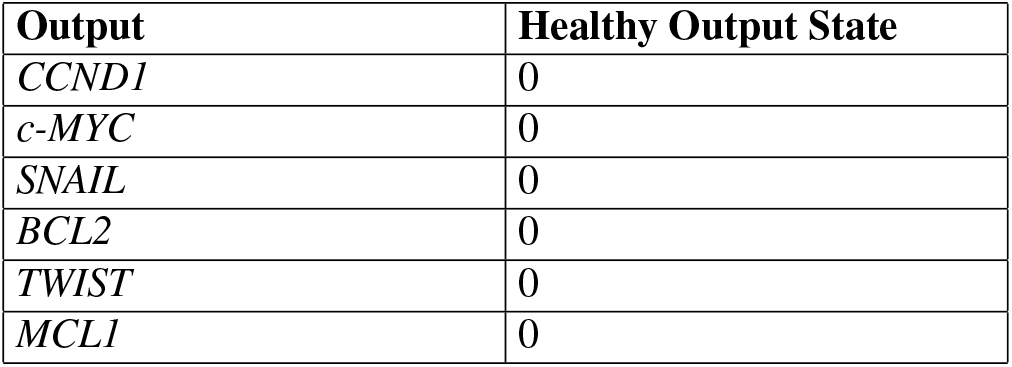
Boolean network (fault free) healthy outputs based on a healthy input

| Output | Healthy Output State |
| --- | --- |
| *CCND1* | 0 |
| *c-MYC* | 0 |
| *SNAIL* | 0 |
| *BCL2* | 0 |
| *TWIST* | 0 |
| *MCL1* | 0 |

When a healthy input signal is applied to the BN in the presence of faults, the output deviates from the healthy output state. Thus applying drugs to a network with faults has the potential to drive the output towards the healthy output. In order to measure the efficacy of each drug, it is necessary to measure how dissimilar the drugged output is from the healthy output state. We quantify the difference between two output vectors using the size difference (SD) score. SD measures the dissimilarity between two binary valued vectors, and its value proportionally increases with a higher dissimilarity.

To mathematically describe SD, suppose we have two binary (0 or 1) valued vectors a= (a_1_,…a_*n*_) and (b_1_,…b_*n*_), where n represents the size of the vectors. Then we can construct a confusion matrix M as follows:

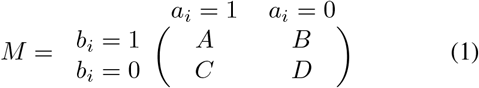

In matrix M, A is the count of cases when the i^*th*^ element of vectors a and b are 1, B is the count of cases when the i^*th*^ element of vector a is 0 and b is 1, C is the count of cases when the i^*th*^ element of vectors a is 1 and b is 0, and D is the count of cases when the i^*th*^ element of vectors a and b are 0,. Thus A and D are the counts of matches between vectors, whereas B and C are counts of mismatches between vectors. Therefore, with the confusion matrix M, we define SD as follows:

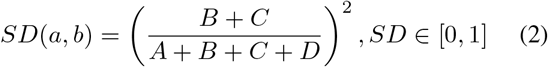

SD ranges from 0 to 1, where a value of 0 indicates both the vectors are identical, and a score of 1 means the highest possible divergence between the vectors (i.e. one vector is a bitwise complement of the other vector). Therefore the higher the SD value the larger the difference between the two vectors. The healthy output state in our fault-free BN requires all the output genes to be in state 0; however, when faults (i.e., mutations) are introduced in this network, the output genes deviate from the healthy output state. Therefore our objective in this study is to identify the drug or combination of drugs that drives the output genes in the presence of faults close to the healthy output state. We can determine this by comparing the drugged output to the healthy output using the SD score. A higher SD value would indicate that the drug is unable to suppress the cancerous output genes in the network, whereas an SD value closer to 0 would imply the drug is effective in suppressing the cancerous output genes. To evaluate drug efficacies, we considered that faults could occur simultaneously in the network, i.e., the TNBCSC pathway can have more than one mutation at a time. We limit our study to a maximum of three faults (mutations) at a time due to computational complexity. Since we have 42 individual faults, we evaluate drug efficacies across a total of ^42^*C*_1_ + ^42^*C*_2_ + ^42^*C*_3_ = 12383 combinations of faults. In addition to the 11 drug classes, which can be administered one at a time, we also wanted to evaluate the results of using a combination of drugs. In our study, we consider a maximum of 5 drug classes that can be applied at a time due to computational complexity. Furthermore, administering too many drugs to a patient at the same time can have adverse effects. For each drug or drug combination, we determine its drugged output across each of the 12383 faults and then compare the drugged output to healthy output and calculate the SD. Thus we will have a resulting matrix containing the SD across each fault for all the considered drugs and their combinations. An example of this resulting matrix of SD is displayed in Table 4, for a network consisting of 3 fault locations. The SD is calculated for three drugs and when no drug is applied to the network in the presence of faults (untreated).

**TABLE 4:** Example table showing SD for three drugs and three faults

|  | Fault 1 | Fault 2 | Fault 3 | Normalized Mean SD |
| --- | --- | --- | --- | --- |
| Drug 1 | 0.2 | 0.8 | 0.6 | 0.666 |
| Drug 2 | 0.3 | 0.5 | 0.2 | 0.416 |
| Drug 3 | 0.4 | 0.2 | 0.1 | 0.292 |
| Untreated | 0.7 | 0.9 | 0.8 | 1.000 |

In Table 4, we have an example matrix of SD for three faults and three drugs. To determine which drug is the most effective for a given fault, we select the drug with the lowest SD. For fault 1, drug 1 has the lowest SD; therefore, it is the most effective drug to suppress fault 1. Similarly, for faults 2 and 3, drug 3 is the best choice. However, to determine the most effective drug across all three faults, we calculate the normalized mean size difference across all the faults. We normalize the mean SD with respect to the mean SD for the untreated case, and this is tabulated in the normalized mean SD column of Table 4. Since drug 3 has the lowest normalized mean SD, it is the best drug to work across all three faults. The normalized mean SD (NMSD) is calculated as follows:

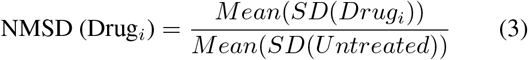

We simulate the BN and calculate the SD using the R programming language [142]. The code for simulating the BN and generating the SD tables are made publicly available at the following GitHub repository: https://github.com/adilahiri/TNBCSC and in the supplemental file section of this paper.

## IV. RESULTS

We calculated the normalized mean SD for each of the drug combinations for the BN with one, two, and three faults (or mutations) at a time. Since there are 11 drug classes and we have considered up to five drugs combinations at a time, this yields a total of ^11^*C*_1_ + ^11^*C*_2_ + ^11^*C*_3_ +^11^*C*_4_ + ^11^*C*_5_ = 1024 drug combinations. With 12383 combinations of faults, our resulting matrix of NMSD would be of size 1024 (drugs) by 12383 (faults). Since it is not tractable to present and comprehend such a large matrix, we have presented the NMSD scores for each drug combination under one, two, and three mutations(or faults) as tables in the supplementary files section of this paper (One_Mutation_File.xlsx, Two_Mutation_File.xlsx,Three_Mutation_File.xlsx). Furthermore, in each supplemental table, we have also presented the top 10 effective drug combinations (i.e., single-drug, two-drug combination,…, and five-drug combination) for each fault combination(these are presented as sub-sheets within each Microsoft Excel file). In the following subsections, we will discuss the ten most effective drug combinations for each fault combination.

### A. DRUG EFFICACY RANKING FOR SINGLE MUTATION NETWORK

We simulated the BN when only one mutation was present at a time for each drug combination scenario. After simulating the BN for each of the 42 faults at a time and calculating the NMSD, we found that the most effective single drug class with the lowest NMSD score was the *NF-κB* inhibitor, followed by *AKT* inhibitor, and *STAT3* inhibitor. Fig 4. displays the drugs with the ten lowest NMSD scores and the untreated case when the network is simulated for one mutation at a time. We observe in Fig.4 (a) that *NF-κB* inhibitor only achieves an NMSD score of 0.5498, we would ideally want this score to be as close to 0 as possible. Thus *NF-κB* inhibitor is not very effective in inhibiting the cancer promoting activities of the output genes. However, compared to the case when no drugs are applied (Untreated) where NMSD is 1, the *NF-κB* inhibitor performs significantly well. Therefore, to drive the NMSD score as close to 0 as possible, we simulate the BN with two, three, four, and five drug combinations at a time. The results of these simulations are shown in Fig.4 (b)-(e), respectively. When two drugs are applied simultaneously, the lowest NMSD score is achieved by the combination of *GLI* and *NF-κB* inhibitors, followed by the combinations of *AKT* and *STAT3* inhibitors. We tabulate the most effective drug combination strategies from a single drug to five drug combinations in Table 5. It is evident from Fig.4 and Table 5 that as the number of drugs applied simultaneously increases, the NMSD score decreases. In Table 5, except for the case when three drugs are applied simultaneously, all the most effective drug intervention strategies include the *NF-κB* inhibitor. For the three drug combination cases, the lowest NMSD score is achieved by the drug combinations of *GLI, AKT*, and *STAT3*, we notice the NMSD score drops approximately by 51% compared to the best single drug intervention strategy consisting of the *NF-κB* inhibitor. Upon increasing the number of the drug combinations, we see the NMSD score (see Table 5) for the most effective drug combination approaches closer to 0, and the most frequently appearing drug inhibitor classes are the *GLI* and *NF-κB* (4 occurrences each), followed by *AKT* and *STAT3* (3 occurrences each). The NMSD score for all the drug combinations for single mutation BN is compiled in Supplemental File 1 (One_Mutation_File.xlsx). Therefore, targeting *GLI* and *NF-κB* is the most effective strategy when there is only one mutation in the network.

**TABLE 5:** Most effective drug intervention strategy for one mutation at time

| Drug Combination | Normalized Mean SD |
| --- | --- |
| *NF-κB* | 0.54985755 |
| *GLI* + *NF-κB* | 0.407407407407407 |
| *GLI* + *AKT* + *STAT3* | 0.267806268 |
| *GLI* + *AKT* + *NFKB* + *STAT3* | 0.165242165242165 |
| *β-Catenin* + *GLI* + *AKT* + *NF-κB* + *STAT3* | 0.0740740740740741 |

**FIGURE 4:**
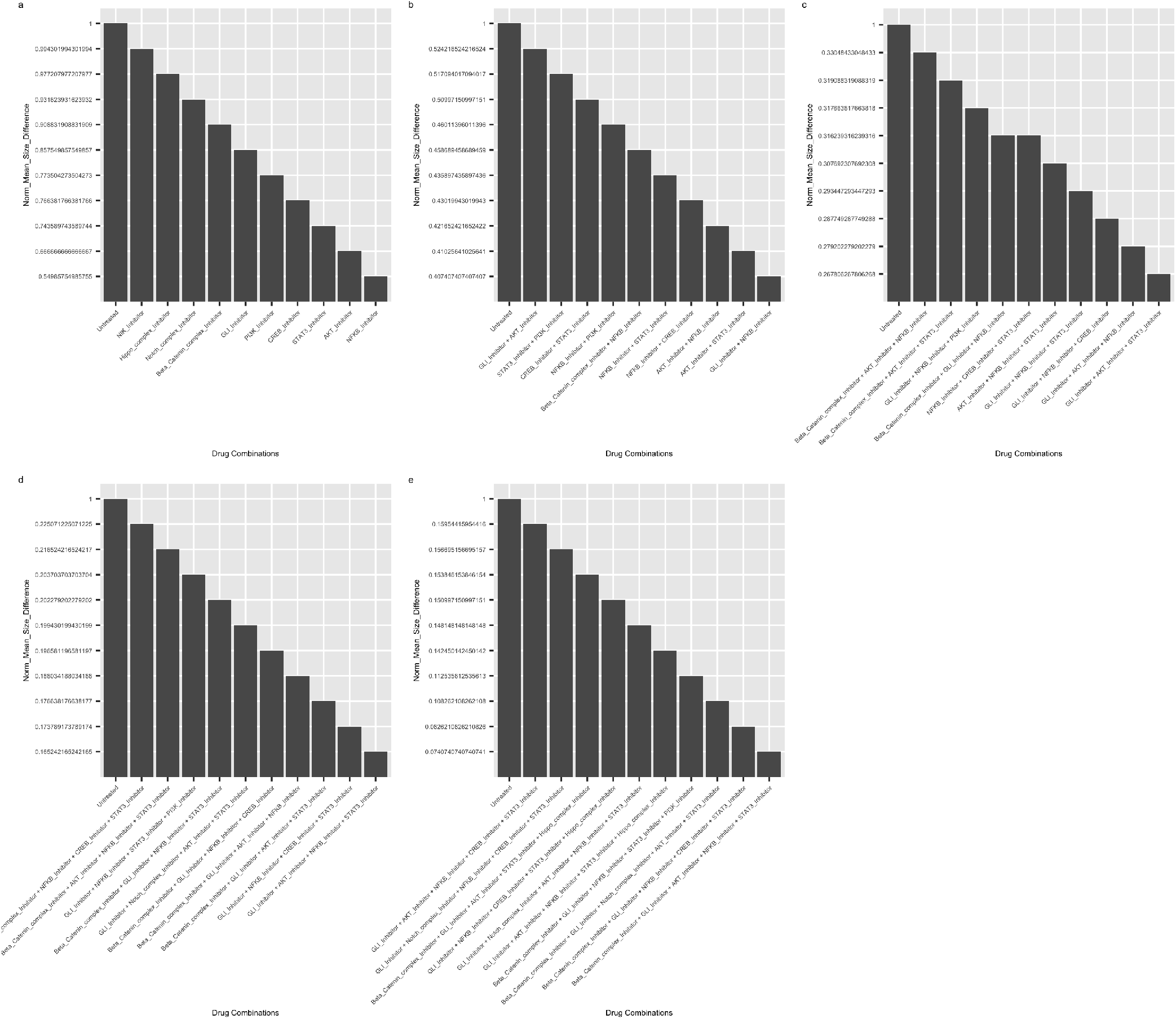
NMSD plots for BN with one mutation at a time. a. NMSD for one drug at a time. b. NMSD for two-drug combinations. c. NMSD for three-drug combinations. d. NMSD for four-drug combination. e. NMSD for five-drug combinations.

### B. DRUG EFFICACY RANKING FOR DOUBLE MUTATION NETWORK

In the previous section, we discussed the drug efficacies when the BN was affected by a single mutation (fault) at a time. However, cancer networks can be afflicted by multiple mutations at a time; therefore, in this section, we simulate the BN with two mutations at a time and calculate the NMSD scores for each drug intervention strategy, as we did in the last section. Fig 5. displays the drugs with the ten lowest NMSD scores and the untreated case when the network is simulated for two mutations at a time. We also tabulate the most effective strategies from one drug to five drug combinations in Table 6. The most effective single drug combination is achieved by the *NF-κB* inhibitor, followed by the *AKT* and *CREB* inhibitors (see Fig.5 (a)). *NF-κB* and *AKT* were also the top two inhibitors when we simulated the network for single mutation in the last section. Unlike the case of single mutations, *STAT3* inhibitor ranked 4th after *CREB* inhibitor under two mutations at a time. The combination of *AKT* and *STAT3* inhibitors is the best two drug combination, followed by the combination of *NF-κB* and *CREB* inhibitors (Fig.5(b)). When three drugs are applied simultaneously, the best combination is *GLI, AKT*, and *STAT3* inhibitors (Fig.5(c)), and it reduces the score difference by 51% compared to the case when only the *NF-κB* inhibitor is applied. This reduction in NMSD is similar to the case of the BN with a single mutation, observed in the prior section. As we increase the number of drugs to four and five, we see the NMSD scores reduce, and the lowest NMSD scores for the most effective drug combination get closer to 0 in both cases (Table 6 and Fig.5 (d,e)). Comparing Tables 5 and 6, and Figs. 4 and 5, we observe that the NMSD score increases as the number of mutations increases in the BN. This is because, with the increased number of mutations, the cancer signals propagate through more pathways, thus the drug combinations need to be able to suppress all the cancer promoting pathways to be effective. Under the two mutations at a time case, we notice that the *STAT3* inhibitor appears the most among the best drug combinations, followed by *GLI* and *AKT* inhibitors. Therefore, targeting *STAT3* seems to be the most effective strategy when there are two mutations in the network. The NMSD score for all the drug combinations for two mutations at a time in the BN are compiled in Supplemental File 2 (Two_Mutation_File.xlsx).

**TABLE 6:**
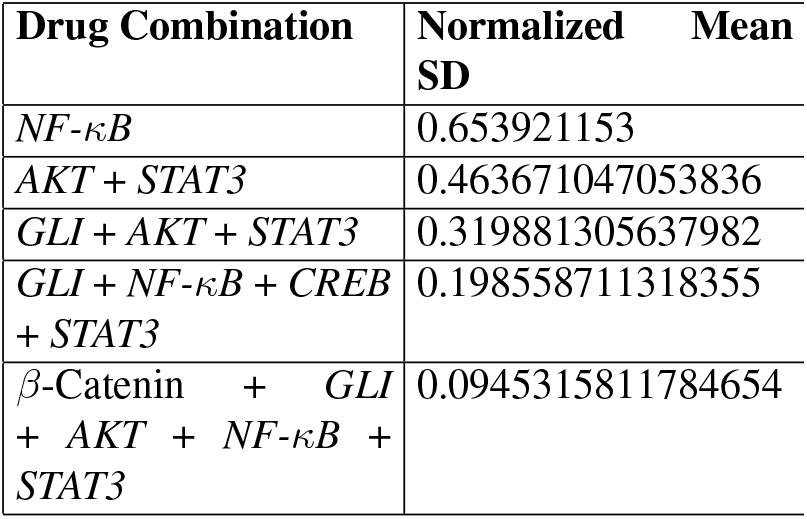
Most effective drug intervention strategy for two mutations at a time

**FIGURE 5:**
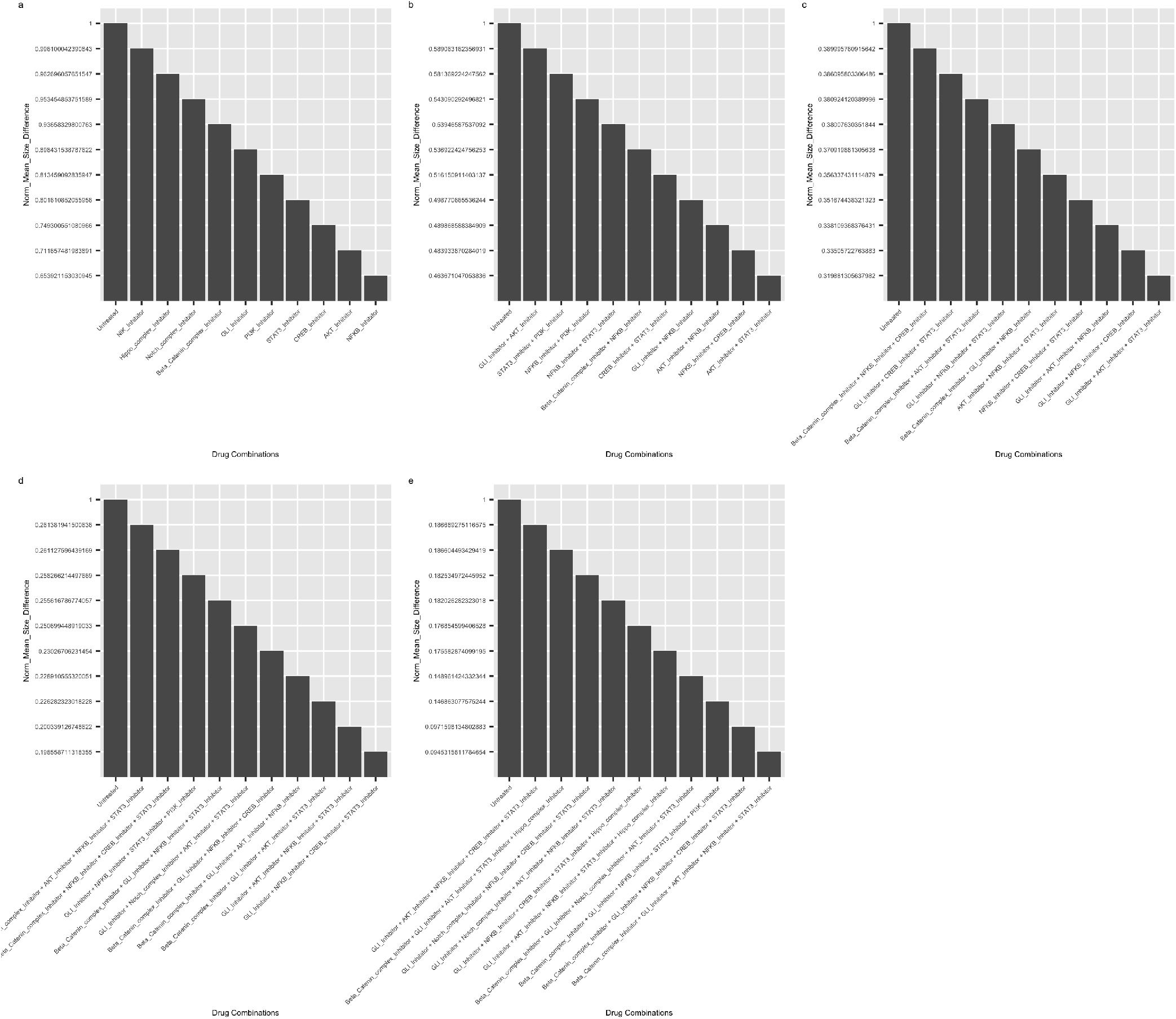
NMSD plots for BN with two mutations a time. a. NMSD for one drug at a time. b. NMSD for two-drug combinations. c. NMSD for three-drug combinations. d. NMSD for four-drug combinations. e. NMSD for five-drug combinations.

### C. DRUG EFFICACY RANKING FOR TRIPLE MUTATION NETWORK

We also simulated the BN for three mutations at a time for each drug combination. We plot the ten most effective intervention strategies from single drug to five drug combinations in Fig 6 and tabulate the top drug combinations from each category in Table 7. From Fig 6 (a), it is clear that *NF-κB* inhibitor is the best single drug intervention, however, its effectiveness under three mutations at a time has reduced compared to two and one mutations at a time case. The combination of *STAT3* and *AKT* inhibitors is the best two drug intervention strategy under three mutations at a time, just like it was for the two mutation case. However, the NMSD score achieved by the combination of *STAT3* and *AKT* inhibitors is still above the halfway mark of 0.5 (Fig.6 (b)), which reflects that even the best two-drug combination is not sufficient in suppressing the cancerous signaling propagated by three simultaneous mutations in the BN. After applying three drugs, we notice that the NMSD score for all ten most effective three-drug combinations falls below 0.5 (Fig.6 (c)). The most effective three-drug combination consists of *GLI, AKT*, and *STAT3* inhibitors, and the NMSD score is approximately 0.37. The only difference between the best two-drug and three-drug combinations is the addition of the *GLI* inhibitor to the latter. By adding *GLI*, the NMSD score drops by approximately 27.89%. As consistent with one and two mutation cases increasing the number of drugs does indeed reduce the NMSD scores. From Table 7, we observe that under three mutations, the *STAT3* inhibitor appears most frequently in the drug combination compared to the other drug inhibitors. It is then followed by *GLI* and *NF-κB* inhibitors. Therefore, targeting *STAT3* is the most effective strategy when there are three mutations in the network. The NMSD score for all the drug combinations for three mutations BN is compiled in Supplemental File 3 (Three_Mutation_File.xlsx).

**TABLE 7:** Most effective drug intervention strategy for three mutations at a time

| Drug Combination | Normalized Mean SD |
| --- | --- |
| *NF-κB* | 0.736607451067486 |
| *AKT + STAT3* | 0.511881798377498 |
| *GLI + AKT + STAT3* | 0.369098923072956 |
| *GLI + NF-κB + CREB + STAT3* | 0.226165492585781 |
| *β-Catenin + GLI + NF-κB + CREB + STAT3* | 0.113546889 |

**FIGURE 6:**
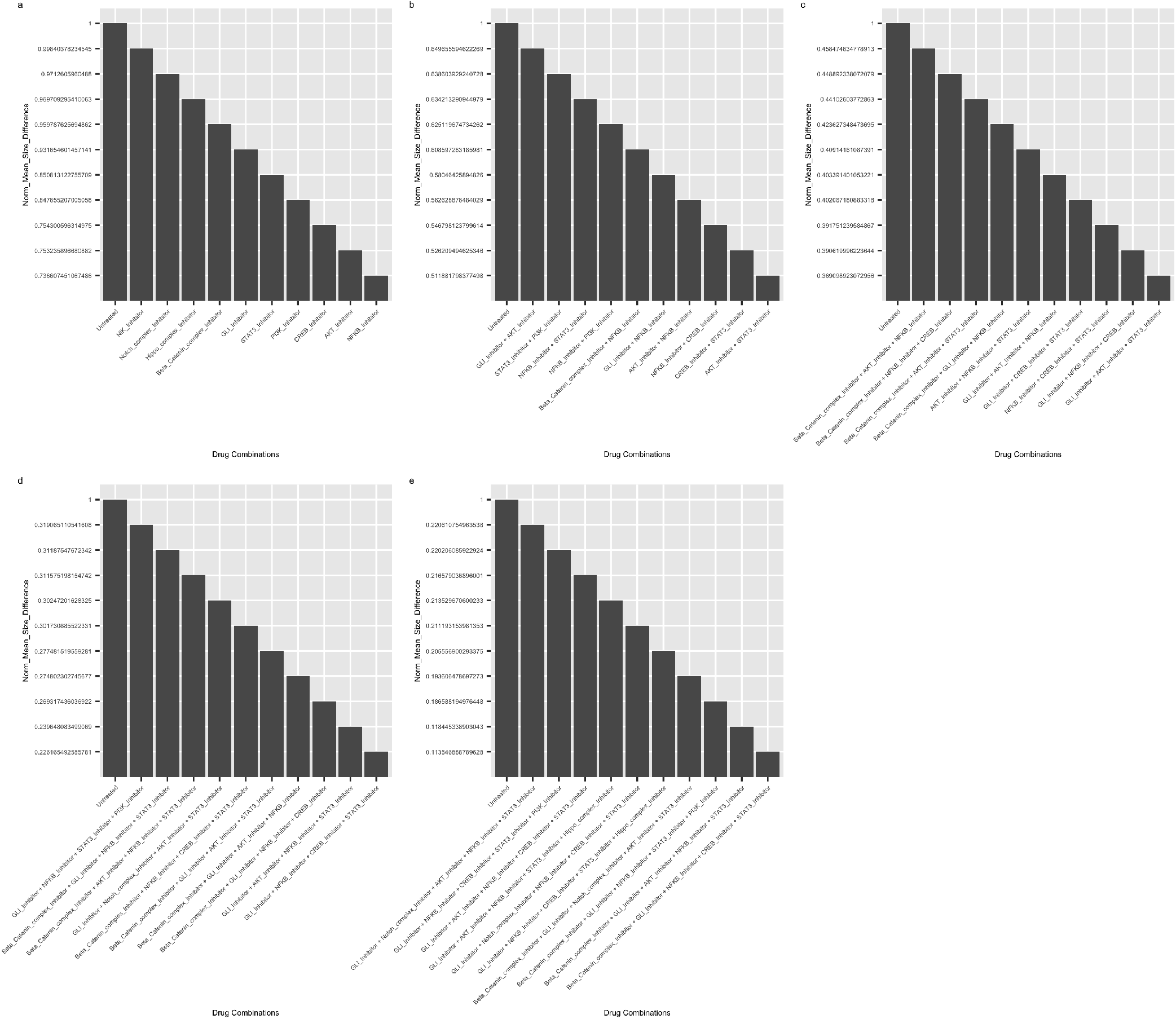
NMSD plots for BN with three mutations a time. a. NMSD for one drug at a time. b. NMSD for two-drug combinations. c. NMSD for three-drug combinations. d. NMSD for four-drug combinations. e. NMSD for five-drug combinations.

## V. DISCUSSION AND FUTURE WORK

In this paper, we created a BN for the TNBCSC signaling pathways from the existing biological literature consisting of both experimental studies and reviews. While BNs are deterministic models, they are an appropriate choice of a modeling paradigm for this study, as publicly available data on TNBCSC are scarce. A search of the NCBI Geo Database using search terms “TNBC Stem Cells” and “Triple Negative Breast Cancer Stem Cells” in February 2023 returned 74 datasets [143]. A majority of the datasets found contained gene expression data for primary tumor cells instead of stem cells. The dataset with the highest number of samples for TNBC stem cells contained only 24 samples, which is insufficient for use in a data-driven model [144]. Thus datadriven or probabilistic modeling was not a reliable choice for this case. One of the drawbacks of the BN is the inability to integrate drug dosage into the model. Drugs often have side effects which need to be considered by clinicians before they are administered to patients. Drug dosage is a critical factor, especially when treating children, the elderly, and patients with comorbidity. In such cases, clinicians might prefer alternative drugs from the same drug class (i.e., an alternative drug that has the same target as the standard of care drug) but with a lower level of dosage and side effects. The dosage information will be useful when comparing two drugs belonging to the same drug class. Since the BN identifies the drug target (i.e. the drug class), the dosage information does not affect our analysis. However, there are computational models that integrate drug dosage and pathway information to predict drug efficiency. Çubuk et al. developed a mechanistic model to simulate the drug interventions and unravel how drug action mechanisms are affected by gender-specific differential gene expression across different cancers [145]. Chen et al. also developed a pathway-integrated model, using mass action modeling to predict the input-output behavior of ErbB signaling pathways, this model can also be further expanded to determine cellular sensitivity to small molecule drugs [146]. These methods of integrating drug dosage are data-driven and when sufficient TNBC stem cell data is available, these methods can serve as a starting point for integrating drug dosage into the TNBCSC signaling pathway. A non-data-driven method that integrated signaling network topology and molecular docking was demonstrated by Gu et al. to evaluate drug efficacy in LPS-induced PGE2 production network [147]. While this model does not require any training data, the authors used their domain knowledge to select proteins for molecular docking and validated the results of their computational methods experimentally. Thus there are several methods that we can consider in our future studies to integrate drug dosage into the TNBCSC signaling pathways.

We simulated the BN with one, two, and three faults or mutations at a time and evaluated the efficacies of various drug combinations in suppressing the six cancer-promoting genes *CCND1, c-MYC, SNAIL,BCL2, MCL1*, and *TWIST*. mWe considered eleven different inhibitory drug classes (Table 1) in our study; these drugs work by binding to their target molecule and inhibiting their downstream signaling. In a mutation-free BN, under a healthy input state, all the cancer-promoting genes were turned off; however, as we modeled mutations into this network, these genes got turned on due to the aberrant signaling from the mutation sites. As the number of turned on cancer-promoting genes increased, the more they deviated from their healthy state, as measured by size difference (SD) score. We applied various drug combinations to the mutated networks to reduce the SD score, the drug combinations with a SD score closer to 0 are considered effective, while drugs with a SD score close to 1 are deemed ineffective. To measure the overall effectiveness of drugs across all combinations of mutations (or faults), we introduced the normalized mean size difference (SD). Similar to the SD metric, an NMSD score close to 1 reflected that the drug was ineffective, and a score close to 0 indicated an effective drug.

From our simulations in section IV, we observed that in the BN with one mutation at a time, we needed at least a two-drug combination to bring the NMSD score below the halfway mark of 0.5. The best two-drug combination was *GLI* and *NF-κB* inhibitors. *GLI* regulates *CCND1, c-MYC*, SNAL, *TWIST, BCL2*, and *NF-κB* regulates *SNAIL* and *TWIST*, thus inhibiting *GLI* and *NF-κB* has the potential to suppress five out of the six cancer-promoting genes in our BN. These five genes promote cell cycle progression, proliferation, and EMT. Only *MCL1* (cell survival), is not directly regulated by *GLI* or *NF-κB*. Thus to drive the NMSD score closer to 0, *MCL1* also needs to be suppressed. In Table 5, for the three, four, and five-drug combinations cases, we notice the addition of *AKT, STAT3*, and *β-Catenin* inhibitors successfully drives the NMSD below 0.5. *AKT* regulates *MCL1* through *CREB, STAT3* directly regulates *MCL1*, thus using drug inhibitors for each of these classes can suppress *MCL1*. While *β-Catenin* does not regulate *MCL1*, it regulates other cancer-promoting genes which reduces the NMSD score down by approximately 55% compared to the four-drug intervention case. We found that *GLI* and *NF-κB* inhibitors appeared the most among the top effective drug interventions (Table 5), when the BN had one mutation. This implies that inhibiting *GLI* and *NF-κB* is essential in TNBCSC networks with one mutation.

We also simulated our BN to include two mutations at a time. The most effective single drug combination for this scenario was again the *NF-κB* inhibitor (same as for the one mutation network). However, compared to the one mutation case, the NMSD score of the *NF-κB* inhibitor increased due to the increased number of mutations in the network. Applying two drugs simultaneously brings the NMSD score below 0.5, and with an increasing number of drugs, the NMSD score gradually decreases. From Table 6, we noticed each of the most effective drug combinations includes *STAT3* (except the case of single drug). *STAT3* directly regulates expressions of the cancer promoting genes *CCND1, c-MYC, MCL1*, and *BCL2*, thus it is highly influential in affecting cancerous activities such as cell cycle progression, cell proliferation, cell survival and apoptosis inhibition. Thus targeting *STAT3* alone, has the potential to suppress all the non-EMT cancer genes in the network and is a primary inhibition target in the two-mutation network.

Our final set of simulations was for the BN consisting of three mutations at a time. Consistent with the one and two mutation cases, the *NF-κB* inhibitor was the most effective single drug for the three mutation case as well. However, the NMSD score for the *NF-κB* inhibitor was approximately 0.74, which is close to the untreated case for which NMSD is Thus with three mutations in the network, applying the *NF-κB* inhibitor by itself or any other drug alone was ineffective in suppressing the cancer-promoting genes. We noticed the NMSD score dropped below 0.5 only when we applied three drugs at a time, and the best three-drug combination consisted of *GLI, AKT*, and *STAT3*. Together, these three genes regulate all six cancer-promoting genes in the BN and are able to control the aberrant cancerous signaling arising from the three mutational sites in the network. However, even with five-drug combinations (Table 7), the lowest NMSD score is approximately 0.11354, which is approximately 53.2 % and 20% higher than the best five-drug combination scores in networks with one and two mutations, respectively. This indicated that increasing the number of drugs is not a very effective strategy, when the number of mutations also increased in the network. We notice that *STAT3* was again the most frequently appearing drug target for the three mutation network, followed by *GLI* and *NF-κB*.

In general, across all our simulations, we noticed that as the number of mutations increased in the network, the NMSD scores also increased, this is because the mutations are able to propagate their cancerous signals through multiple pathways and evade the inhibitory effects of the drugs. We observed across our simulations that increasing the number of drugs applied does decrease the NMSD score, but with increasing number of mutations, this strategy is not effective and is also not practical as a patient cannot be given too many drugs at the same time. Based on our simulations, we noticed that if the network has more than one mutation, targeting *STAT3* is essential, followed by *NF-κB*, and *GLI*. When applying only one drug, the *NF-κB* inhibitor was the best single drug across the networks containing one, two, and three mutations.

*STAT3,NF-κB*, and *GLI* regulate all the six cancer-promoting genes in the BN and are thus the best targets for inhibition. In contrast, drugs that inhibit upstream targets, such as *NIK* inhibitor, *PI3K* inhibitor, are not as effective. Drugs that suppress upstream targets are not effective in suppressing cancerous signaling when mutations lie downstream from their target, whereas drugs that affect downstream targets are mostly able to counteract upstream mutations. Furthermore multiple studies have indicated that *STAT3,NF-κB*, and *GLI* are potential drug targets for TNBC [115], [148]–[150]. Thus we can conclude that inhibiting *STAT3, NF-κB*, and *GLI* is a potentially essential strategy for suppressing cancerous signaling in the TNBCSC pathways.

The next steps for this study would involve validating our findings by carrying out wet lab experiments on cell lines or on patient-derived tumor xenograft models (PDX) and evaluating the effect of *STAT3, NF-κB*, and *GLI* inhibitors in lowering the expression levels of the six cancer-promoting genes in the network. A lowered expression of these genes would be a preliminary indicator of the efficacy of drugs and would warrant testing the drugs on higher-order animal models and eventually reaching the preclinical trial stage.

Furthermore, we want to explore the landscape of mutations in gene regulatory networks and evaluate the impact it has on drug development and drug repurposing with an overarching goal of enhancing personalized medicine. Knowledge of the degree to which a mutation or a group of mutations drive aberrant gene regulatory behavior also has implications for clinical trial design, including mutation-focused basket-trial formats that are becoming increasingly popular in the era of precision medicine [151], [152].

A possible application for the current BN model could be to help formulate inclusion criteria for basket-trials by specifying not only the type of mutations but also the number of mutations permitted in the study. Basket trials are a type of clinical trials where multiple targeted therapies are tested in patients who have different types of cancer that all have the same mutations or biomarkers [153]. Our results in this study showed that with an increasing number of mutations, even the most optimal single-drug intervention is not effective. Furthermore, we demonstrated that a combination of multiple drugs is able to suppress the aberrant signaling from multiple mutation sites. Thus, our model can potentially be used to determine the maximum number and type of mutations under which a drug or combinations of drugs might be effective and accordingly define the eligible patient pool for basket trials.

## VI. CONCLUSION

TNBC is an aggressive form of breast cancer with a high rate of relapse. While chemotherapy, which is the standard of care treatment for TNBC is effective in targeting the primary tumor cells, it often fails to target TNBC stem cells which are responsible for relapse. In this study, we modeled the underlying biological pathways in TNBC stem cells using BNs and evaluated the efficacy of small molecule inhibitory drugs in suppressing cell cycle progression, proliferation, survival, apoptosis inhibition, and EMT. Our findings indicated that combinations of drugs are better at suppressing cancerous activity compared to single drugs, especially in the presence of multiple mutations in the pathways. We found that targeting and inhibiting *STAT3, NF-κB*, and *GLI* is an essential strategy in suppressing cancerous activity in TNBC stem cells. Therefore, *STAT3 NF-κB*, and *GLI* are potential drug targets for TNBC stem cells and prevent relapse. Applying targeted therapies for treating TNBC will be a novel treatment methodology, as it is not part of the standard of care for the treatment of TNBC which currently includes only chemotherapy. Unlike chemotherapy, targeted therapies have fewer adverse side effects which is particularly critical when the patients are older adults. Therefore, if targeted therapies can prevent a TNBC patient from relapsing and undergoing a second cycle of chemotherapy, it will significantly improve the quality of life of the patient. However, it must be noted that chemotherapy is still the standard of care and is necessary for treating the primary tumor. Applying targeted therapies which include inhibitors of *STAT3 NF-κB*, and *GLI* alongside chemotherapy has the potential to target both the TNBC stem cells and primary tumor cells and prevent future relapse and could become a new treatment paradigm for combating TNBC.

